# GYS2 Promotes the Progression of Aortic Dissection and Aortic Aneurysm via C5a/NF-κB-Mediated Pro-Inflammatory Macrophage Polarization

**DOI:** 10.64898/2026.06.08.731008

**Authors:** Lei Jin, Likang Ma, Juncheng Liu, Zongsheng Jiang, Wangting Wu, Yuanxin Liu, Anqi Xie, Xueping Huang, Xinmeng Xie, Liangwan Chen, Li Zhang, Zhihuang Qiu

## Abstract

**Background:** Aortic aneurysm and aortic dissection (AAD) are lethal cardiovascular emergencies characterized by sudden onset and extremely high early mortality. The pro-inflammatory polarization of macrophages is one of the core factors driving the pathogenesis of AAD, but its underlying mechanism remains unclear. This study focuses on the aberrant expression of glycogen synthase 2 (GYS2) in the AAD microenvironment and its role in driving macrophage polarization toward the pro-inflammatory M1 phenotype.

**Methods:** Proteomic analysis was conducted to identify protein heterogeneity associated with AAD. Clinical and animal samples were used to evaluate the correlation between GYS2 expression and AAD progression. Whole-body GYS2 knockout mice and adeno-associated virus (AAV)-mediated macrophage-specific gain- and loss-of-function models were utilized to investigate the regulatory role of GYS2 in macrophage polarization and complement activation. Downstream molecular pathways were identified and validated through in vitro stimulation and in vivo exogenous C5a rescue experiments.

**Results:** GYS2 expression was significantly upregulated in AAD tissues and primarily localized in macrophages. Activation of GYS2 by LiCl or macrophage-specific overexpression of GYS2 exacerbated aortic dilation and extracellular matrix degradation, and increased mortality in AAD mice. Conversely, whole-body GYS2 knockout or macrophage-specific GYS2 knockdown suppressed inflammatory factors, significantly reduced the incidence of AAD, and attenuated vascular injury. Mechanistically, excessive GYS2 in macrophages specifically triggered the complement-coagulation cascade, promoting the generation of the potent anaphylatoxin C5a. C5a further bound to its receptor C5AR1, activating the downstream PLCβ3/NF-κB signaling pathway, thereby inducing M1 macrophage polarization and matrix metalloproteinase-mediated extracellular matrix degradation. In vivo exogenous C5a rescue completely reversed the vascular protective effects conferred by GYS2 deficiency.

**Conclusions:** This study demonstrates that highly expressed GYS2 regulates the pro-inflammatory polarization of macrophages and extracellular matrix degradation via the complement C5a/PLCβ3/NF-κB signaling axis, which is a key mechanism driving AAD progression. Specific inhibition of macrophage GYS2 can effectively alleviate aortic vascular inflammation and prevent AAD progression, providing a promising novel strategy for the clinical conservative treatment of AAD.

## INTRODUCTION

Aortic dissection or aortic aneurysm (AAD) is a sudden-onset, catastrophic cardiovascular emergency [1, 2]. Its early mortality rate is extremely high, and the risk of death increases progressively within the first few hours after symptom onset [3]. According to the latest Global Burden of Disease (GBD) 2021 report, approximately 153,927 deaths were attributed to aortic dissection and aneurysm, marking a 74.2% increase since 1990 and posing a formidable challenge to current clinical management [4, 5]. The typical pathological feature of AAD is that following an intimal tear, high-pressure blood flow directly impacts the aortic media, causing extensive dissection of the medial layer and forming a true and false double-lumen structure[6]. Currently, there is a lack of effective drugs to prevent the structural degradation of the aortic wall or reverse the underlying pathological progression, and emergency surgical intervention remains the only life-saving strategy. Therefore, elucidating the core molecular mechanisms of AAD pathogenesis and identifying key therapeutic targets for local vascular deterioration is urgently needed.

To identify this key upstream regulatory molecule, we conducted high-throughput proteomic screening of aortic tissues from diseased mice and, for the first time, identified that glycogen synthase 2 (GYS2) is aberrantly overexpressed in AAD lesions. As a classical rate-limiting enzyme in the glycogen synthesis pathway [15], GYS2 has historically been studied primarily for its role in maintaining systemic glucose homeostasis [16]. However, emerging research in immunometabolism has demonstrated that the aberrant expression of metabolic enzymes can directly modulate immune cell phenotypes through metabolic reprogramming, acting as critical signaling hubs in chronic vascular inflammation [17, 18]. Although studies have suggested that excessive activation of the complement and coagulation cascade is a significant contributor to aortic wall inflammatory remodeling [19, 20], the precise pathophysiological function of GYS2 in cardiovascular diseases, particularly within the AAD inflammatory microenvironment and macrophage polarization, remains unexplored.

In this study, we demonstrated via in vivo animal models that the aberrant overexpression of GYS2 significantly exacerbates β-aminopropionitrile (BAPN)-induced aortic dissection in mice. Conversely, both global GYS2 knockout and macrophage-specific deletion via AAV-mediated gene engineering effectively reduced the incidence of AAD and the severity of vascular structural destruction. In-depth mechanistic investigations revealed that GYS2 accumulation within macrophages specifically activates the complement coagulation cascade, promoting the generation of the potent pro-inflammatory anaphylatoxin C5a. This, in turn, promotes macrophage polarization toward the M1 phenotype and excessive extracellular matrix degradation via the C5a/NF-κB signaling pathway. This study not only fills the research gap regarding the key role of GYS2 in the regulation of vascular immunometabolism, but also provides a promising molecular strategy for the development of targeted, non-surgical interventions for aortic dissection or aortic aneurysm.

## Methods

### Human Tissue Samples and Ethics Approval

The 20 aortic dissection (AAD) specimens included in this study were all obtained from patients undergoing open repair surgery, and the diagnosis of all cases was confirmed by senior cardiothoracic surgeons based on intraoperative exploration and postoperative histopathological evaluation. The control group vascular specimens (n=15) were derived from organ donors or non-AAD patients undergoing heart transplantation, and their detailed clinical baseline data are shown in Table S1. All experimental procedures involving human specimens strictly adhered to the ethical principles of the Declaration of Helsinki, and were reviewed and formally approved in advance by the Ethics Committee of Fujian Medical University Union Hospital (Ethics approval number: 2026KY461). To protect patient privacy, all clinical data and specimen information were strictly de-identified and encrypted for storage, and the security management of the data was executed in full compliance with the biomedical research guidelines of the institution.

### Animal Studies and Ethics Approval

All animal experiments were conducted in strict compliance with the Guide for the Care and Use of Laboratory Animals (8th edition, 2011) published by the NIH [21] and approved by the Animal Care and Use Committee of the Institute of Laboratory Animal Science, Fujian Medical University (Approval No.: 2026-Y-1368). The experiments were performed in accordance with the “3Rs” principles[22]. All mice were housed in a standardized specific pathogen-free (SPF) facility under a 12/12 h light/dark cycle at 26±2°C and 50%±5% humidity, with ad libitum access to standard chow and sterile water. Interventional experiments were initiated after a 1-week acclimatization period. Given that clinical epidemiological data indicate a significantly higher incidence of AAD in males, only male mice were used to avoid the confounding effects of sex hormones [23].

### Source of Mice and Genetic Models

Wild-type (WT) C57BL/6J male mice (1 or 3 weeks old) were purchased from Wu’s Laboratory Animal Technology Co., Ltd. GYS2⁻/⁻ mice were generated by Cyagen Biosciences; genotypes were confirmed by PCR. For macrophage-specific overexpression, mice were injected via the retro-orbital vein with 2×10¹¹ vg/mouse of rAAV9-F4/80-GYS2-FLAG. For macrophage-specific knockdown, mice were treated with rAAV9-F4/80-sgGYS2 at the same titer, with rAAV9-Null serving as the control.

### BAPN-induced AAD Mouse Model

The AAD model was established using 0.25% (w/v) BAPN in drinking water for 28 days. Subsequently, Ang II (GLPBIO, USA) was administered via intraperitoneal injection (1.44 mg/kg, three times daily for 3 days). At the experimental endpoint, mice were deeply anesthetized with 1% pentobarbital sodium for surgery and tissue collection.

### Cell Culture and Drug Treatment

Cells were maintained in DMEM with 10% FBS at 37°C in a 5% CO₂ incubator. Before drug treatment, cells were serum-starved for 6 hours. Cells were then treated with Ang II (Tocris, UK), LPS (MedChem Express, USA), or recombinant C5a (1 μg/mL, MedChem Express, USA) for the indicated durations.

### Western Blotting

Protein was extracted using RIPA buffer and quantified via BCA assay. After SDS-PAGE and transfer to PVDF membranes, membranes were incubated with primary antibodies against GYS2, IL-1β, MMP2, MMP9, P65, p-P65, IκBα, p-IκBα, PLCβ3, p-PLCβ3, C5, C5AR1, and GAPDH. After incubation with HRP-conjugated secondary antibodies, signals were detected using ECL reagents and quantified via ImageJ.

### Quantitative Real-time PCR (qRT-PCR)

Total RNA was extracted using TRIzol reagent and reverse-transcribed. qPCR was performed using an Applied Biosystems 7900HT system. GAPDH served as the internal control. Primer sequences are provided in Table S2.

### Immunofluorescence (IF) and Immunohistochemistry (IHC)

Paraffin sections (4 μm) were deparaffinized, rehydrated, and subjected to antigen retrieval. For IF, sections were blocked with 5% BSA and incubated with primary antibodies, followed by fluorescent secondary antibodies and DAPI. For IHC, HRP-conjugated secondary antibodies were used, followed by DAB visualization and hematoxylin counterstaining. Images were captured using a Pannoramic MIDI scanner and confocal microscopy.

### Cellular Immunofluorescence

Cells seeded in confocal dishes were fixed with 4% paraformaldehyde, permeabilized with 0.1% Triton X-100, and incubated with primary/secondary antibodies and DAPI. Fluorescence intensity was quantified using ImageJ.

### Histological Staining

HE, EVG (Solarbio, China), and Masson’s Trichrome (Solarbio, China) staining were performed according to the manufacturers’ instructions.

### Flow Cytometry and Cell Sorting

For phenotypic analysis, cells were stained with fluorescent antibodies or DCFH-DA (Solarbio, China) and analyzed on a FACSVerse flow cytometer. For macrophage sorting, single-cell suspensions from aortic tissue were stained with F4/80-APC and CD11b-PE and isolated using a FACS Aria sorter.

### ELISA

Target cytokine (TNF-α, IL-6) and C5a levels were measured in serum and cell supernatants using commercial ELISA kits.

### Small Animal Ultrasound

At day 28, mice were anesthetized with isoflurane. Aortic diameter and blood flow were measured using a high-frequency ultrasound system (Vevo series). Measurements were performed by two independent, blinded researchers.

### Statistical Analysis

Data were analyzed using GraphPad Prism 9, SPSS 23.0, and R 4.1.0. Normality was assessed via the Shapiro-Wilk test, and variances were compared using the F-test. Data are expressed as mean ± SD. Two-group comparisons were performed using the unpaired Student’s t-test (normal) or Mann-Whitney U test (non-normal). Multiple group comparisons were performed via one-way/two-way ANOVA with Bonferroni post hoc tests or the Kruskal-Wallis test with Dunn’s multiple comparisons. Survival analysis was performed using Kaplan-Meier curves and the log-rank test. P < 0.05 was considered statistically significant.

## Results

### GYS2 is significantly upregulated in the vascular lesion microenvironment of AAD patients and mice

To investigate the core drivers of aortic dissection (AAD), we performed quantitative proteomics on aortic tissues from β-aminopropionitrile (BAPN)-induced AAD mice and PBS-treated controls. Principal component analysis (PCA) revealed distinct separation between the two groups in global protein expression profiles (Figure 1A). Subsequent differential expression analysis precisely identified dysregulated targets in AAD vascular lesions (Figure 1B), and hierarchical clustering heatmaps highlighted the distinct molecular signatures between the two groups (Figure 1C). Utilizing the LASSO regression algorithm, we identified four potential biomarkers with diagnostic value, and their predictive efficacy was confirmed via ROC curves (Figure S1A–1C). Notably, Glycogen synthase 2 (GYS2) exhibited a highly significant upregulation in AAD tissues (Figure 1D).

**Figure 1.**
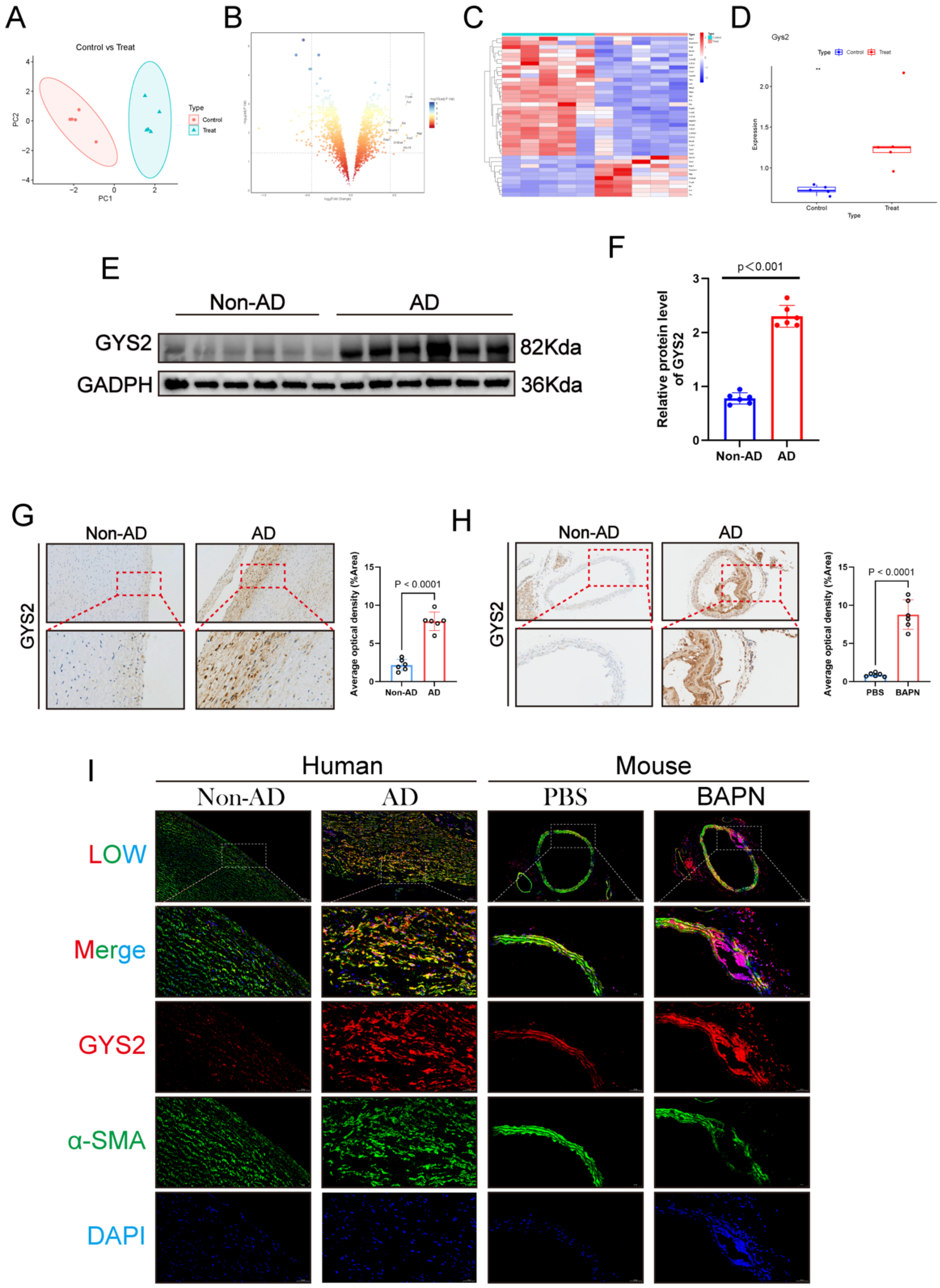
GYS2 is upregulated in human and mouse AAD. **A:** Principal component analysis (PCA) plot of differentially expressed proteins from the proteomics data. **B:** Volcano plot and **C:** hierarchical clustering heatmap of the differentially expressed proteins. **D:** Relative expression levels of GYS2 extracted from the proteomics data. **E:** Representative Western blot images of GYS2 protein in human aortic tissues from non-AD and AD patients, and in mouse aortic tissues treated with BAPN or PBS (control). **F:** Quantitative analysis of GYS2 protein expression in human and mouse aortic tissues (*n* = 6). Data were analyzed by Student’s *t*-test. Representative immunohistochemistry (IHC) images and quantitative analysis of mean optical density for GYS2 in **G:** human aortic tissues (non-AD vs. AD) and **H:** mouse aortic tissues (PBS vs. BAPN) (*n* = 6). Data were analyzed by Student’s *t*-test. **I:** Representative double immunofluorescence staining images of GYS2, α-SMA, and DAPI (4′,6-diamidino-2-phenylindole) in human and mouse aortic tissues.

To corroborate our proteomic results, we quantified GYS2 levels in both human and murine AAD specimens. Both Western blotting and immunohistochemistry consistently revealed a marked upregulation of GYS2 protein in human AAD tissues (Figure1E, 1F, 1G) and mouse AAD aortas (Figure 1H). Notably, immunohistochemical analysis showed that GYS2 was highly enriched at the media-adventitia junction, suggesting its potential localization within smooth muscle cells or local immune cell infiltrates. Furthermore, dual immunofluorescence staining for GYS2 and the smooth muscle marker α-SMA (Figure 1I) demonstrated that the aberrantly elevated GYS2 was broadly dispersed throughout the adventitia in both human and mouse aortas. This localization strongly indicates that adventitia-infiltrating macrophages serve as the primary cell subset for GYS2-mediated pathological regulation.

### Pharmacological activation of GYS2 exacerbates AAD progression and vascular rupture in mice

To elucidate the role of GYS2 activation in the progression of aortic dissection (AAD), we used lithium chloride (LiCl, a classic GYS activator) for pharmacological intervention in a BAPN-induced mouse model of AAD [24, 25]. A total of 120 four-week-old male C57 mice were randomly allocated into four equal groups: control (water + PBS), LiCl, BAPN, and LiCl+BAPN (Figure 2A). Baseline body weights showed no significant fluctuations across all groups throughout the experiment (Figure 2B). Following Ang II injection on day 28, mortality in the BAPN group was 16.6% (n=5), whereas this rate surged to 30% (n=9) in the LiCl+BAPN group. Statistical data revealed that 56.6% (n=17) of mice in both of these groups developed an aortic dissection or aneurysm (AAD) (Figures 2C and 2D). Aortic ultrasound, however, demonstrated that LiCl intervention significantly worsened BAPN-induced aortic dilation (Figure 2E). Histological staining (H&E, Masson’s trichrome, and EVG) further confirmed that GYS2 activation severely aggravated vascular matrix degradation and pathological collagen deposition (Figure 2F). In contrast, the LiCl group showed no obvious vascular toxicity from LiCl alone (Figure 2F). Collectively, these results demonstrate that GYS2 activation exacerbates the formation, progression, and rupture of AAD.

**Figure 2.**
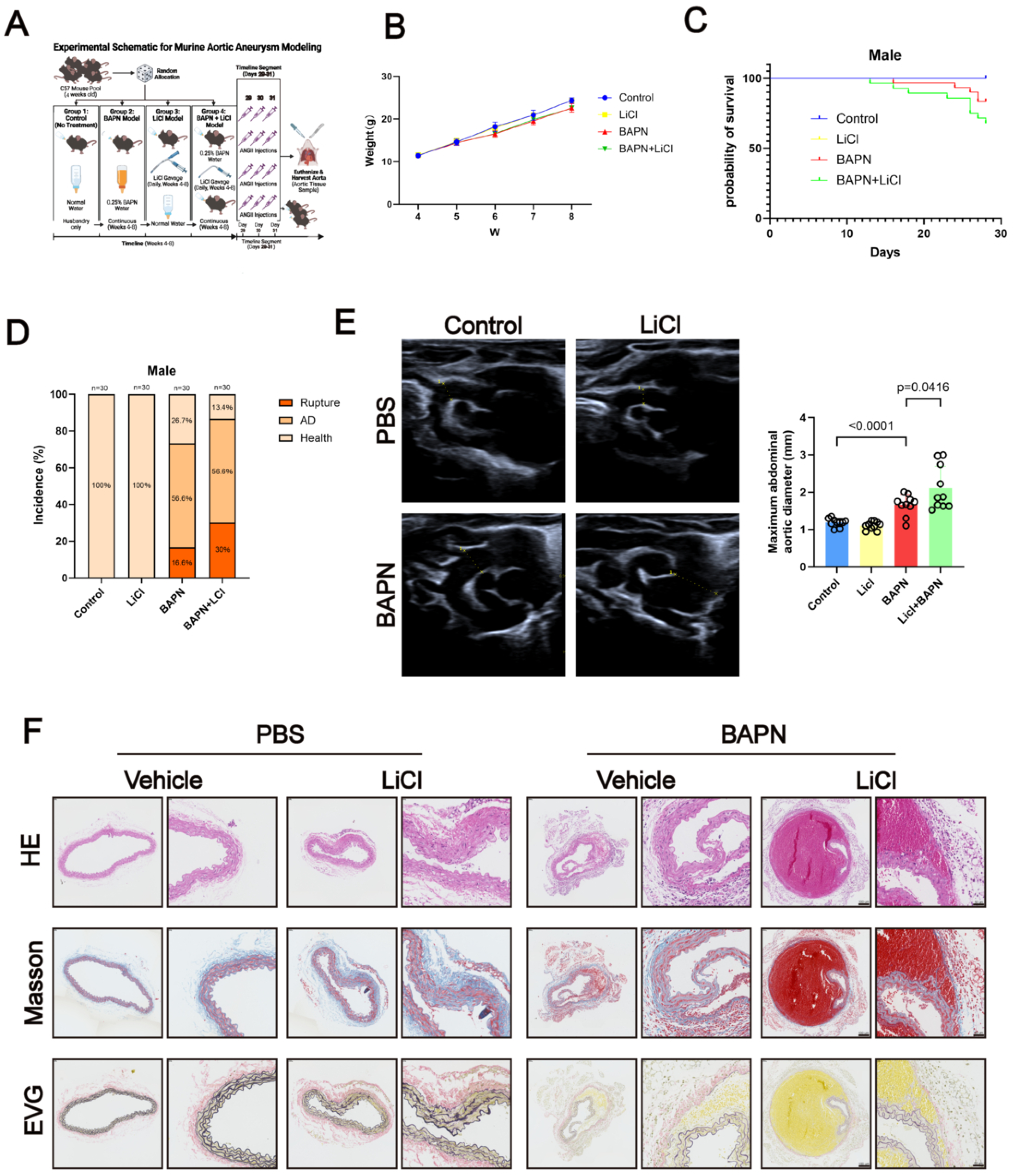
LiCl exacerbates BAPN-induced mouse aortic dissection formation and rupture. **A:** Schematic diagram of the experimental model of mouse aortic dissection. **B:** Body weight changes of the designated groups during the modeling period. **C:** Kaplan-Meier survival curves of male mice. **D:** Incidence of aortic complications, including rupture, dissection (AD), and health (*n* = 30 per group). **E:** Representative ultrasound images of the thoracic aorta (*n* = 10]per group). **F:** Representative hematoxylin-eosin (H&E), Masson, and elastica van Gieson (EVG) staining images of the aorta. Data are expressed as mean ± SD. Statistical analysis of diameter measurements was performed by one-way ANOVA followed by Tukey’s *post hoc* test, and survival rates were analyzed by Log-rank test.

### Global GYS2 deficiency suppresses AAD progression and ameliorates local inflammatory remodeling

To determine the direct target effect of GYS2 in AAD injury, we generated GYS2 knockout (GYS2⁻/⁻) mice. Four groups were established: control, GYS2⁻/⁻, BAPN, and GYS2⁻/⁻+BAPN (Figure 3A). Physical signs remained stable (Figure 3B). After 28 days, the BAPN group mortality reached 23.3% (n=7), whereas global GYS2 deficiency reduced the mortality to 10% (n=3). Simultaneously, the AAD incidence in the GYS2⁻/⁻+BAPN group (40%, n=12) was significantly lower than in the BAPN group (53.4%, n=16) (Figure 3C, 3D). Ultrasound confirmed that GYS2 knockout effectively suppressed aortic wall dilation (Figure 3E, 3F). Multi-histological staining (HE, Masson, EVG) revealed that GYS2 deficiency greatly attenuated elastic lamellar fragmentation and collagen accumulation (Figure 3G).

**Figure 3.**
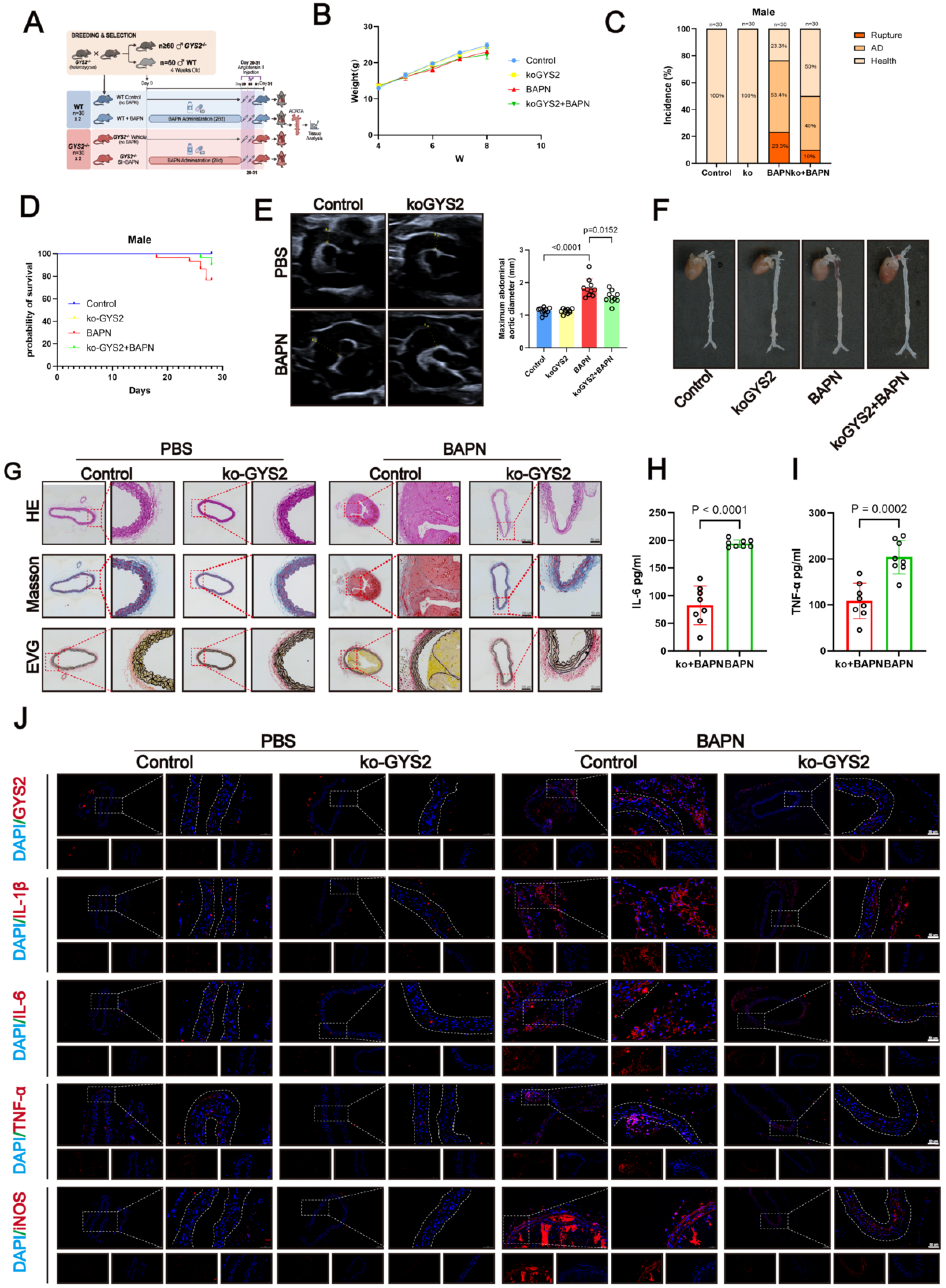
Global deletion of GYS2 attenuates BAPN-induced mouse aortic dissection formation and inflammatory response. **A:** Schematic diagram of the experimental design. Four-week-old male wild-type (WT) and global GYS2 knockout (GYS2−/−) mice were treated with or without BAPN for 28 days (Control, GYS2−/− Vehicle, BAPN, and ko+BAPN groups, *n* = 30 per group). **B:** Body weight fluctuations of mice in each group during the experimental period. **C:** Incidence of aortic complications (rupture, AD, and health) in the designated animal cohorts. **D:** Survival curves of the same animal cohorts as in (A). **E:** Representative ultrasound images of the abdominal aorta and measurements of the maximal aortic diameter on day 28. **F:** Representative gross images of the aorta from the experimental cohorts. **G:** Representative histological analysis of aortic sections, including H&E, Masson’s trichrome, and EVG staining. **H-I:** Quantitative analysis of serum IL-6 **(H)** and TNF-α **(I)** levels measured by ELISA. **J:** Representative immunofluorescence images of GYS2, IL-1β, IL-6, TNF-α, and iNOS in aortic cross-sections. Nuclei were counterstained with DAPI. All data are presented as mean ± SD. Statistical analysis of the maximal aortic diameter **(E)** and ELISA results **(H-I)** was performed using one-way ANOVA. Survival curves **(D)** were evaluated using the Log-rank test.

Given the central role of inflammation, we tracked peripheral inflammatory markers. ELISA confirmed that plasma IL-6 and TNF-α concentrations in the GYS2⁻/⁻+BAPN group were significantly reduced compared with the BAPN group (Figure 3H, 3I). Immunofluorescence and immunohistochemistry of aortic sections corroborated these results: GYS2 knockout potently quelled the BAPN-induced inflammatory cytokine storm within the vascular wall (Figure 3J and Figure S2A). These data indicate that GYS2 inhibition not only preserves vascular structure but also intercepts pathological inflammatory remodeling at the source.

### Macrophage-specific GYS2 knockdown achieves precise therapeutic rescue of AAD vascular injury

Based on the evidence of GYS2 enrichment in adventitial immune cells, we constructed a macrophage-targeted GYS2 knockdown model using AAV9-F4/80-sgGYS2 (Figure 4A). Flow cytometry sorting and RT-qPCR confirmed GYS2 silencing in the target subpopulation (Figure S3A). Notably, macrophage-specific GYS2 deletion showed superior efficacy over global knockout: the mortality rate in AAV9-sgGYS2 mice approached baseline (3.3%, n=1), and the AAD incidence decreased to 30% (n=10) (Figure 4B, 4C). Ultrasound showed that targeted intervention almost completely blocked aneurysmal dilation (Figure 4D, 4E). Pathological staining confirmed that macrophage GYS2 knockdown preserved elastic fiber integrity and prevented vascular structural collapse (Figure 4F). IF and IHC further confirmed that targeted depletion of GYS2 in macrophages precisely and potently dissolved the local vascular wall inflammatory microenvironment (Figure 4G and Figure S3B).

**Figure 4.**
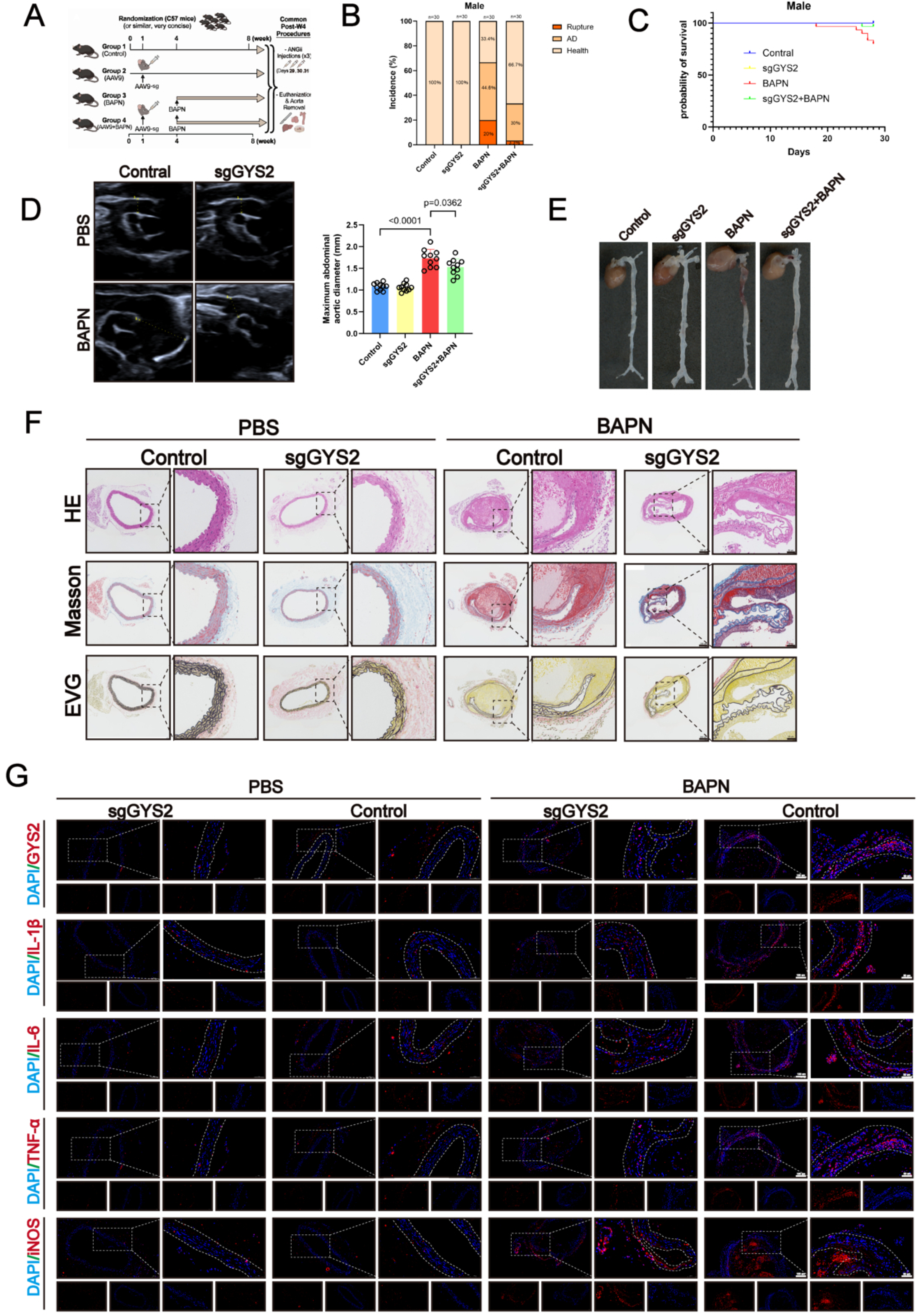
Macrophage-specific knockdown of GYS2 attenuates BAPN-induced mouse aortic dissection and elastin degradation. **A:** Schematic diagram of the experimental design. Two-week-old male C57BL/6 mice received retro-orbital vein injection of AAV9-F4/80-Cas9-sgGys2 to achieve macrophage-specific GYS2 deletion, followed by 28 days of BAPN treatment (*n* = 30 per group). **B-C:** Incidence of aortic complications (B) and Kaplan-Meier survival curves (C) in the designated experimental groups. **D:** Representative ultrasound images of the abdominal aorta and measurements of the maximal aortic diameter on day 28. **E:** Representative gross images of the aorta from the experimental cohorts. **F:** Representative histological analysis of aortic sections, including H&E, Masson’s trichrome, and EVG staining, showing the alleviation of vascular pathology and elastin disorganization. **G:** Representative immunofluorescence images demonstrating the attenuated inflammatory activation in the aortic wall following macrophage-specific GYS2 deletion. All data are presented as mean ± SD. Statistical analysis was performed using one-way ANOVA (E) or Log-rank test (C).

### Macrophage-specific GYS2 overexpression exacerbates local inflammatory storms and matrix collapse

To provide rigorous mechanistic counter-evidence, we utilized AAV9-F4/80-GYS2-FLAG for targeted overexpression in mouse macrophages. Flow cytometry and RT-qPCR verified a significant increase in GYS2 transcriptional levels (Figure S4A). Under BAPN induction (Figure 5A), GYS2 overexpression led to severe deterioration in survival, with mortality surging to 50% (n=15) compared with the BAPN control group (20%, n=6). Furthermore, the AAD incidence in the overexpression group was significantly increased (Figure 5B, 5D), and ultrasound recorded severe aortic dilation near rupture (Figure 5C, 5E). Histologically, GYS2 overexpression acted as a catalyst, disintegrating the continuity of elastic lamellae and inducing catastrophic collagen degradation and matrix collapse (Figure 5F). ELISA and pathological staining confirmed that macrophage GYS2 overexpression directly triggered systemic inflammatory activation (Figure 5G) and local vascular inflammatory cascades (Figure 5H and Figure S4B).

**Figure 5.**
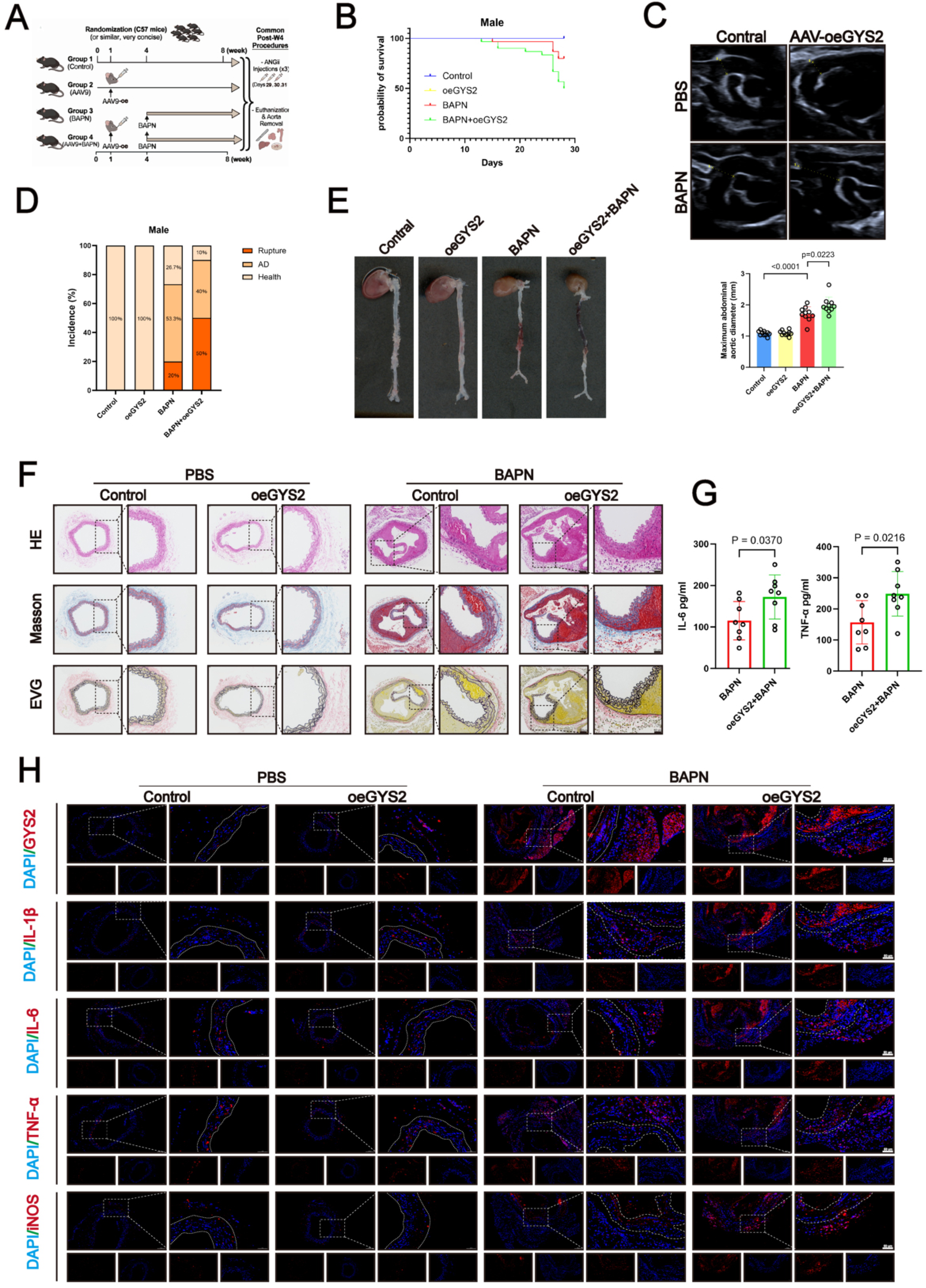
Macrophage-specific GYS2 overexpression exacerbates BAPN-induced mouse aortic dissection and inflammatory response. **A:** Schematic diagram of the experimental design. Two-week-old male C57BL/6 mice received retro-orbital vein injection of AAV9-F4/80-oeGYS2 to achieve macrophage-specific GYS2 overexpression (or control vector), followed by treatment with or without BAPN for 28 days (Control, oeGYS2 Vehicle, BAPN, and AAV-oeGYS2+BAPN groups, *n* = 30 per group). **B, D:** Kaplan-Meier survival curves (B) and incidence of aortic complications (including rupture, AD, and health) (D) in the designated experimental cohorts. **C, E:** Representative ultrasound images of the abdominal aorta on day 28 (C), and representative gross images of the aorta and measurements of the maximal aortic diameter (E). **F:** Representative histological analysis of aortic sections, including H&E, Masson’s trichrome, and EVG staining, showing exacerbated vascular pathology, elastin disorganization, and collagen degradation. **G:** Quantitative analysis of serum IL-6 and TNF-α levels measured by ELISA in the BAPN and AAV-oeGYS2+BAPN groups (*n* = [8] per group). **H:** Representative immunofluorescence images of GYS2, IL-1β, IL-6, TNF-α, and iNOS in aortic cross-sections, highlighting enhanced inflammatory activation. Nuclei were counterstained with DAPI. All data are presented as mean ± SD. Statistical analysis was performed using one-way ANOVA (E), unpaired Student’s *t*-test (G), or Log-rank test (B).

### GYS2 promotes macrophage M1 polarization and induces oxidative stress decompensation

In the AAD lesion core, infiltrating macrophages polarize toward the pro-inflammatory M1 phenotype and release excessive inflammatory mediators and matrix metalloproteinases, serving as core drivers of aortic medial elastic matrix destruction [26]. To explore the function of GYS2, we utilized lipopolysaccharide (LPS) to simulate the AAD microenvironment in vitro [27] and implemented lentiviral GYS2 intervention. Immunofluorescence analysis showed that, under LPS stimulation, GYS2 overexpression significantly amplified the translation levels of M1 polarization markers (iNOS) and core inflammatory molecules (TNF-α, IL-1β, IL-6) (Figure 6B); conversely, GYS2 knockdown suppressed M1 polarization and dampened these molecular signals (Figure 6A).

**Figure 6.**
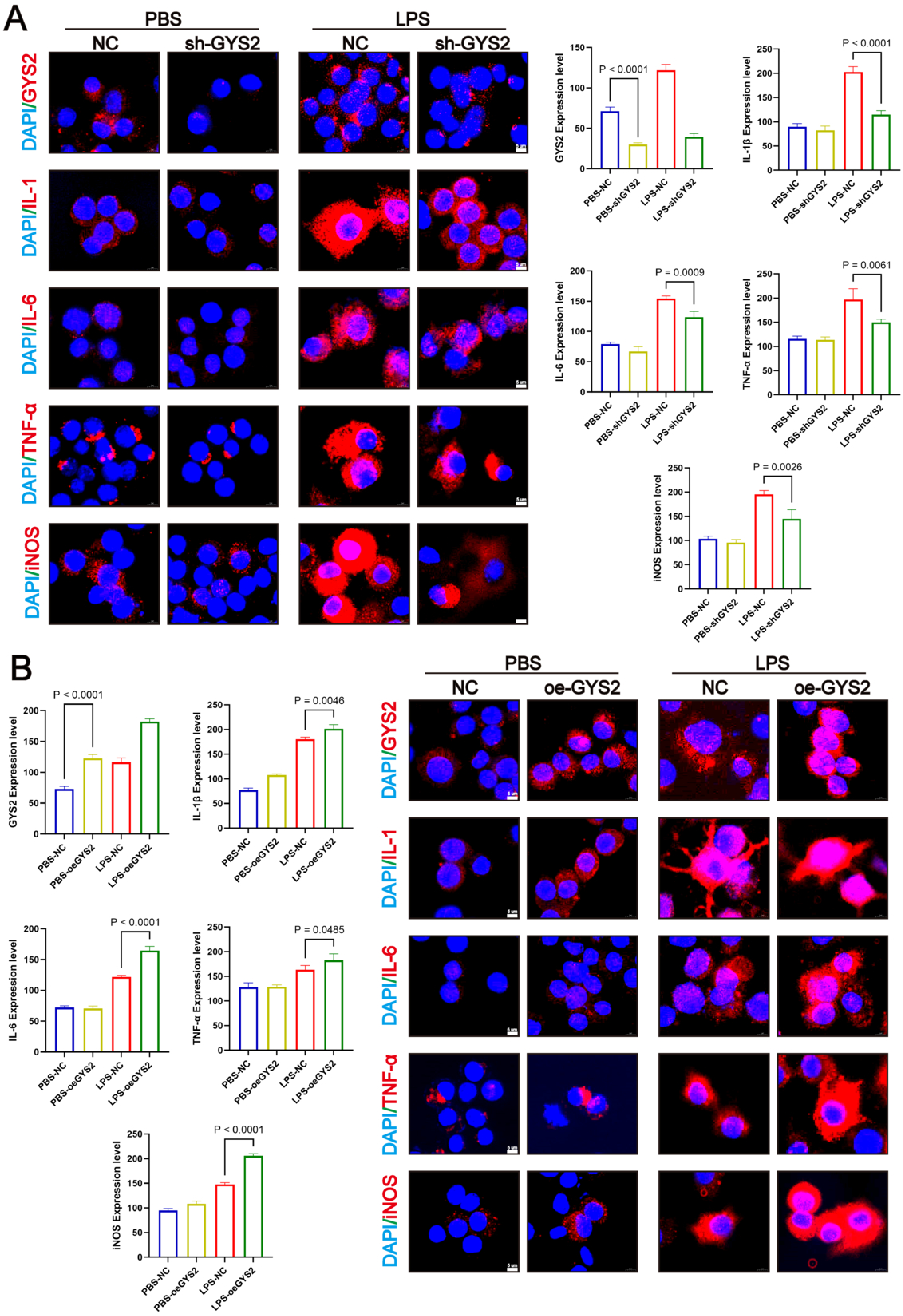
GYS2 drives M1 macrophage polarization and pro-inflammatory cytokine production *in vitro*. **A:** Representative immunofluorescence images and quantitative analysis of GYS2, IL-1β, IL-6, TNF-α, and iNOS expression in macrophages transfected with empty vector control (NC) or sh-GYS2 lentivirus under PBS or LPS (lipopolysaccharide) stimulation. **B:** Quantitative analysis and representative immunofluorescence images of GYS2, IL-1β, IL-6, TNF-α, and iNOS expression in macrophages transfected with empty vector control (NC) or oe-GYS2 lentivirus under PBS or LPS stimulation. Scale bar = 5 µm. Data are expressed as mean ± SD. Statistical analysis was performed by one-way ANOVA followed by Tukey’s *post hoc* test.

Given that robust polarization is often accompanied by redox imbalance [28], we assessed intracellular reactive oxygen species (ROS) via flow cytometry. Results revealed that under pathological stress, GYS2 overexpression directly induced explosive accumulation of ROS in macrophages, pushing oxidative stress into an irreversible decompensation phase (Figure S5A). These experiments confirm that GYS2 is a critical intracellular engine driving M1 polarization and oxidative damage.

### Macrophage GYS2 accelerates extracellular matrix degradation by hijacking the complement-coagulation cascade

To further explore the downstream transcriptional regulatory network of GYS2-induced matrix collapse, we performed high-throughput proteomics on aortic tissues from GYS2-overexpressing mice. Differential expression clustering (FC > 1.2, P < 0.05) captured 146 upregulated and 47 downregulated targets (Figure S6A, S6B). GO analysis strongly indicated that GYS2 abnormalities are highly intertwined with blood coagulation, inflammatory activation, and macromolecular transcriptional regulators (Figure 7A). KEGG enrichment was particularly definitive: the “complement and coagulation cascades” dominated, followed by “ECM-receptor interaction” (Figure 7B). In vitro experiments validated these findings. Lentiviral-mediated GYS2 overexpression under LPS stimulation triggered an explosive increase in the fluorescence intensity of the elastin-degrading enzymes MMP2 and MMP9 (Figure 7D). This direct biochemical evidence suggests that GYS2 hijacks and amplifies the complement cascade, ultimately realizing the pathological effect of large-scale extracellular matrix degradation.

**Figure 7.**
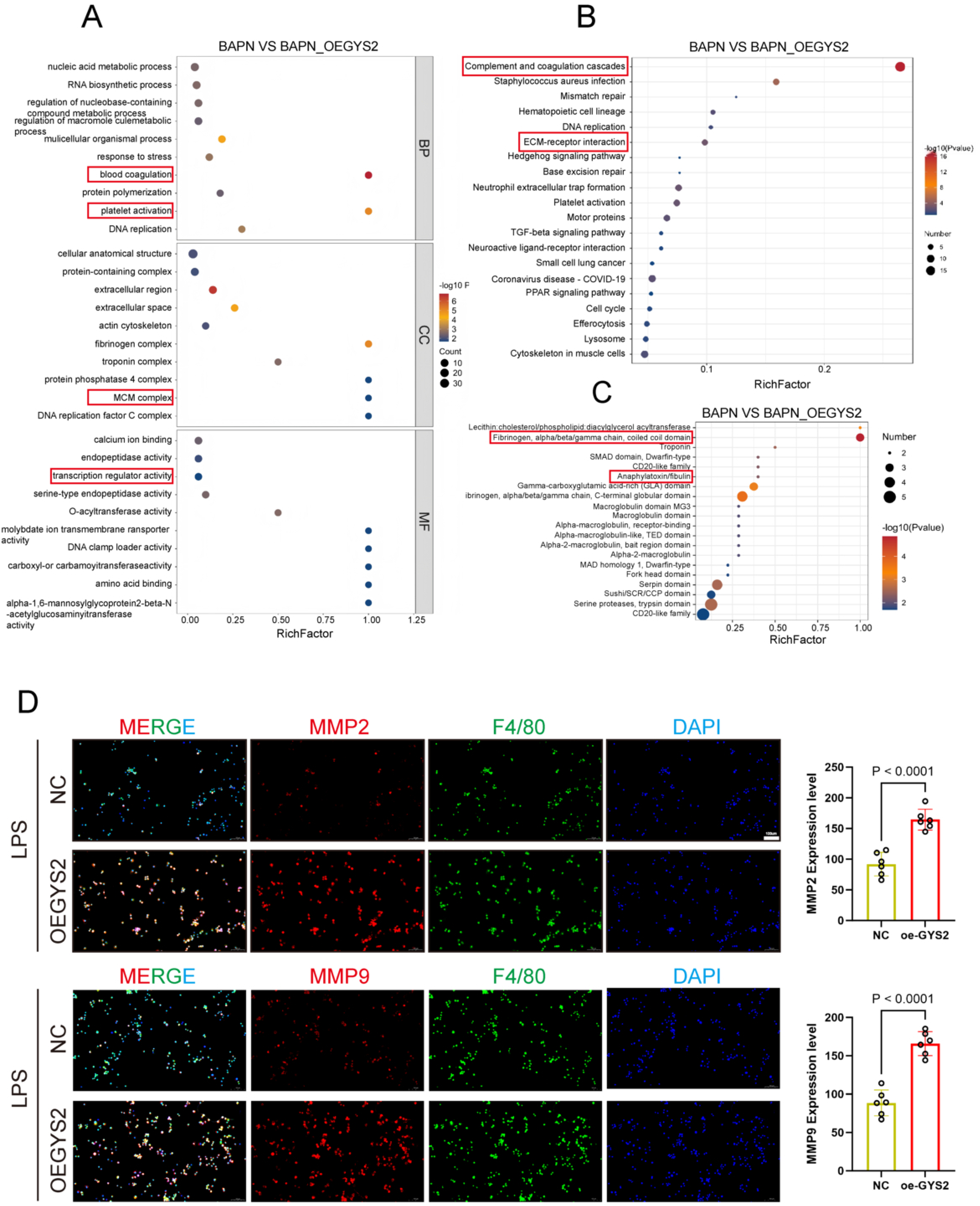
Macrophage GYS2 aggravates aortic dissection through complement and coagulation cascades and extracellular matrix degradation. **A– C:** Proteomic sequencing analysis of mouse aortic tissues from the BAPN and BAPN+AAV-oeGYS2 groups. **(A)** Gene Ontology (GO) enrichment analysis of differentially expressed proteins based on proteomic sequencing of aortic tissues from the two groups of mice (BAPN group and BAPN+oeGYS2 group). Differentially regulated proteins (Fold change > 1.2 and *P* < 0.05) were enriched in biological processes, cellular components, and molecular functions, including blood coagulation, platelet activation, and transcription regulator activity. **(B)** The most significantly changed canonical pathways in the KEGG pathway analysis predicted based on the differentially regulated proteins (*P* < 0.05), revealing that the complement and coagulation cascades and ECM-receptor interactions are the core signaling pathways. **(C)** Protein domain enrichment analysis showing the enrichment of fibrinogen and anaphylatoxin domains.**D:**Representative immunofluorescence images and quantitative analysis showing the co-staining of MMP2 and MMP9 (shown in red) with the macrophage marker F4/80 (shown in green) in macrophages transfected with negative control (NC) or oe-GYS2 lentivirus and stimulated with LPS (*n* = 6 per group). Nuclei were counterstained with DAPI (shown in blue). All data are presented as mean ± SD. Evaluated by Student’s *t*-test (D). BAPN: β-aminopropionitrile; oeGYS2: GYS2 overexpression; GO: Gene Ontology; KEGG: Kyoto Encyclopedia of Genes and Genomes; ECM: extracellular matrix; MMP: matrix metalloproteinase; LPS: lipopolysaccharide; DAPI: 4′,6-diamidino-2-phenylindole.

### GYS2 amplifies local macrophage inflammatory effects via the complement C5a/NF-κB axis

To identify the central hub molecule in the complement system, we performed InterPro domain screening on high-abundance proteins. Our analysis corroborated complement system activation while accurately detecting a widespread enrichment of the Anaphylatoxin/fibulin domain (Figure 7C). This domain acts as the central engine propelling the complement cascade, with specific molecules harboring it including C3a, C4a, and C5a[29, 30]. Proteomic details further confirmed significant upregulation of the upstream hub protein C5. Canonical molecular pathology dictates that C5a, the potent anaphylatoxin generated by C5 cleavage, anchors to its macrophage receptor C5AR1 [31–33], subsequently activating the PLCβ3/NF-κB signaling network via phosphorylation [34–36]. Accordingly, kinase phosphorylation assays in GYS2-overexpressing macrophages confirmed that the phosphorylation sites of p-P65, p-IκBα, and p-PLCβ3 were significantly activated (Figure 8A-8C), and confocal microscopy visually captured massive nuclear translocation of P65 (Figure 8D).

**Figure 8.**
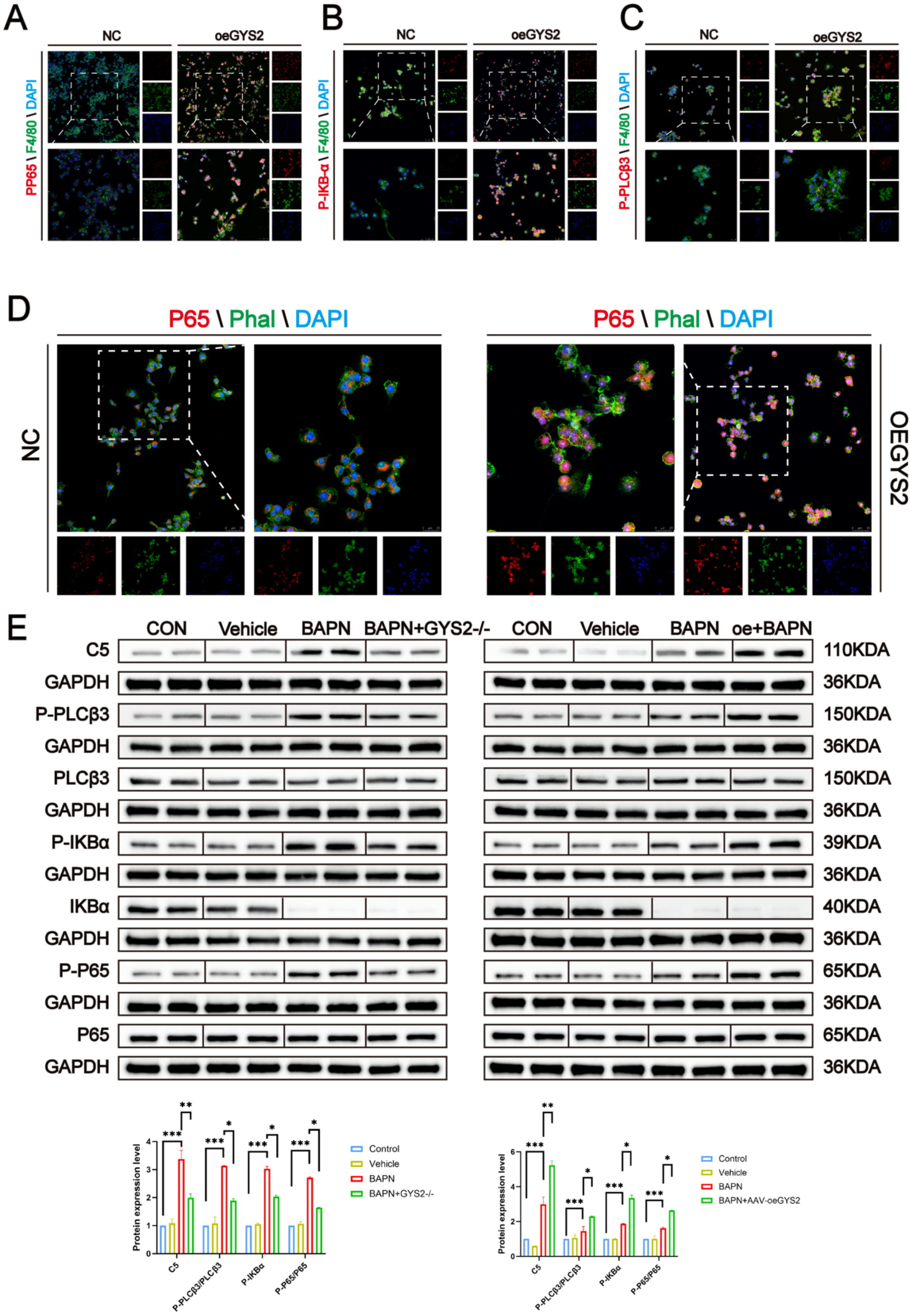
Macrophage GYS2 aggravates macrophage inflammation through the C5-PLCβ3-NF-κB signaling pathway. **A-C**: Representative immunofluorescence images showing the co-staining expression levels of phosphorylated (P)-P65 **(A)**, P-IKBα **(B)**, and P-PLCβ3 **(C)** (shown in red) with the macrophage marker F4/80 (shown in green) in macrophages transfected with negative control (NC) or oeGYS2 lentivirus. Nuclei were counterstained with DAPI (shown in blue). **D:** Representative immunofluorescence images demonstrating the nuclear translocation of P65 (shown in red) in macrophages transfected with NC or oeGYS2 lentivirus. The cytoskeleton was stained with phalloidin (Phal, shown in green), and nuclei were stained with DAPI (shown in blue). **E:** Representative Western blot images and corresponding quantitative analysis of C5, P-PLCβ3, PLCβ3, P-IKBα, IKBα, P-P65, and P65 protein expression in aortic tissues from *in vivo* animal cohorts. The left panel shows the global GYS2 knockout model, and the right panel shows the macrophage-specific GYS2 overexpression model, and their respective control groups. GAPDH was used as an internal control. All data are presented as mean ± SD. Evaluated by one-way ANOVA (E). NC: negative control; oeGYS2: GYS2 overexpression; BAPN: β-aminopropionitrile; DAPI: 4′,6-diamidino-2-phenylindole; Phal: phalloidin; GAPDH: glyceraldehyde-3-phosphate dehydrogenase.

In vivo validation also anchored this signaling axis. In the plasma of AAV-GYS2-overexpressing mice, C5a concentrations surged (Figure S6A); conversely, global GYS2 knockout suppressed circulating C5a (Figure S6B). Combined with Western blot validation in the same two animal models (Figure 8E), this study establishes a rigorous molecular evidence chain: uncontrolled GYS2 in macrophages completes its AAD-aggravating process by relying on the complement-inflammation hub pathway of C5a/PLCβ3/NF-κB.

### Exogenous C5a fully reverses the vascular protective phenotype conferred by GYS2 deficiency

To verify whether C5a is the requisite hub for GYS2-mediated pathological effects, we designed an in vivo “rescue” strategy. Under BAPN induction, we continuously infused exogenous recombinant C5a into GYS2⁻/⁻ mice. The sharp phenotypic reversal confirmed our hypothesis: under identical stress, C5a intervention abolished the AAD-protective effect of GYS2 deficiency, leading to an upward trend in AAD incidence and mouse mortality (Figure 9A, 9B). Real-time aortic diameter monitoring showed that the vascular dilation previously clamped by GYS2 knockout became uncontrolled again (Figure 9C, 9D). Multi-histological sections observed irreversible aortic expansion and severe elastic fiber fragmentation, which re-emerged in the rescue cohort (Figure 9E).

**Figure 9.**
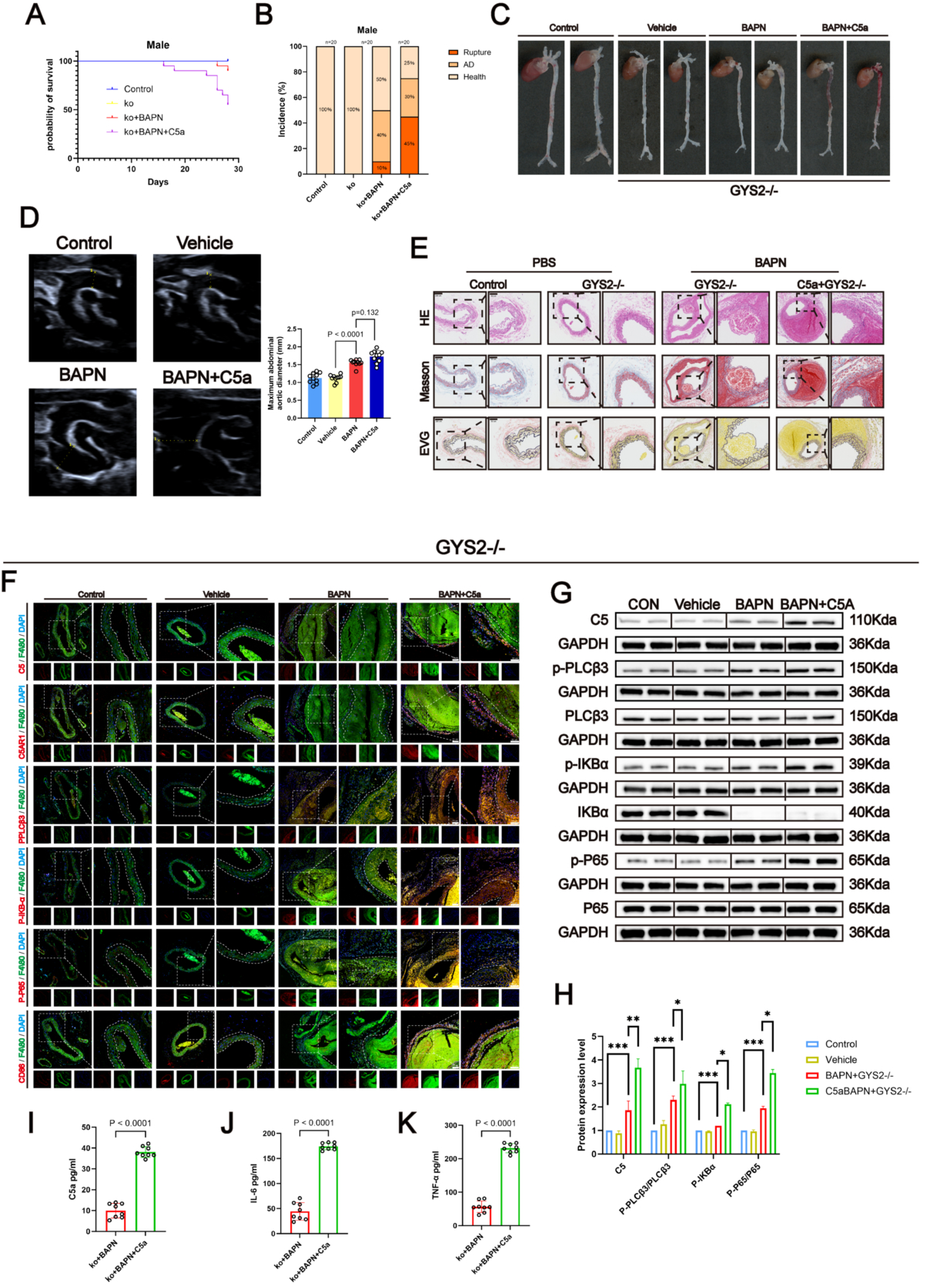
Exogenous C5a supplementation reverses the protective effect of GYS2 deletion against BAPN-induced aortic dissection. **A, B:** Kaplan-Meier survival curves **(A)** and the incidence of aortic complications (including rupture, AD, and health) **(B)** in WT control, GYS2−/− (ko), BAPN-treated ko (ko+BAPN), and C5a-supplemented ko (ko+BAPN+C5a) mice. **C:** Representative gross images of the aortas on day 28, and **D:** representative thoracic aorta ultrasound images and measurements of the maximal aortic diameter. **E:** Representative histological analysis of aortic sections, including H&E, Masson’s trichrome, and EVG staining, indicating that C5a supplementation re-induces vascular pathology and elastin degradation. **F:** Representative immunofluorescence images showing the co-staining of C5, C5AR1, p-PLCβ3, p-IKBα, p-P65, and the M1 macrophage marker CD86 (shown in red) with the macrophage marker F4/80 (shown in green) in aortic sections. Nuclei were counterstained with DAPI (shown in blue). **G:** Representative Western blot images and **H:** corresponding quantitative analysis of C5, p-PLCβ3, PLCβ3, p-IKBα, IKBα, p-P65, and P65 protein expression in aortic tissues. GAPDH was used as an internal control. **I-K:** Quantitative analysis of plasma C5a **(I)**, IL-6 **(J)**, and TNF-α **(K)** levels measured by ELISA. All data are presented as mean ± SD. Evaluated by one-way ANOVA (D, H) or unpaired Student’s *t*-test (I, J, K). AD: aortic dissection; BAPN: β-aminopropionitrile; ko: knockout; EVG: Elastica van Gieson; DAPI: 4′,6-diamidino-2-phenylindole; GAPDH: glyceraldehyde-3-phosphate dehydrogenase; ELISA: enzyme-linked immunosorbent assay.

Molecular data were consistent with the macroscopic phenotype. In situ immunofluorescence (Figure 9F) and Western blotting (Figure 9G, 9H) confirmed that exogenous C5a restored the NF-κB core pathway in the aorta (inducing high p-IκBα and p-P65 expression), awakened downstream receptors C5AR1 and p-PLCβ3, and was accompanied by macrophage re-infiltration. ELISA also captured a rebound in IL-6, TNF-α, and C5a concentrations (Figure 9I-9K). This decisive in vivo rescue experiment confirms: GYS2 in macrophages relies entirely on the cascade amplification of the C5a/NF-κB signaling axis to ultimately drive AAD toward catastrophic rupture.

## Discussion

Exploring the immunometabolic imbalance within large vessels during the progression of acute aortic dissection (AAD) holds significant translational value for overcoming the bottleneck of current treatment strategies that rely heavily on surgical intervention. Based on quantitative proteomics, this study analyzed the complex changes in AAD; a key finding was the aberrant overexpression of glycogen synthase 2 (GYS2) in the vascular remodeling microenvironment, which was highly enriched in locally infiltrated adventitial macrophages. Through rigorous in vivo animal experiments (encompassing both global GYS2 knockout and AAV-mediated macrophage-specific intervention), we constructed a robust chain of evidence: GYS2 abnormality is not merely a biochemical accompaniment to AAD pathogenesis but a core metabolic engine directly driving aortic wall matrix collapse and accelerating lethal rupture. Targeted inhibition of macrophage GYS2 potently quells pathological vascular dilation and matrix degradation, providing a novel perspective for understanding large-vessel immune remodeling.

The non-canonical “immunometabolism-complement” regulatory cascade established in this study provides a solid molecular foundation for deciphering the activation network of pro-inflammatory macrophage polarization (M1). Comprehensive in vivo and in vitro evidence indicates that the pathological accumulation of GYS2, a rate-limiting enzyme within macrophages, acts as the initiating factor for C5 upregulation and the release of the potent pro-inflammatory anaphylatoxin C5a. Through the downstream C5a-C5AR1-PLCβ3-NF-κB cascade, this axis activates and sustains the persistent inflammatory state of macrophages within the vascular wall. This phenomenon, wherein a key enzyme of the glycogen synthesis pathway triggers complement cascade activation, reflects the inherent evolutionary logic of immunometabolism in the AAD lesion microenvironment. From the perspective of intracellular biochemical flux, the sharp changes in the GYS2 expression profile not only interfere with traditional glycogen storage but also reshape metabolism, breaking the original metabolic equilibrium within macrophages. Metabolic reprogramming leads to the disintegration of macrophage homeostasis and localized microenvironmental imbalance [28], thereby precisely driving C5 translation enhancement and protein abundance accumulation at the post-transcriptional level. The surge in C5 protein levels provides an abundant supply of complement substrate precursors to the diseased large vessels, which are catalytically cleaved by the local protease system to explosively generate the highly active anaphylatoxin C5a [29, 30]. The excessive C5a then anchors with high affinity to the specific receptor C5AR1 on the macrophage surface [31–33], triggering precise downward transmission of intracellular kinase signals and potently inducing pathological phosphorylation of the key node PLCβ3 (p-PLCβ3) [34]. Activated p-PLCβ3 efficiently initiates the phosphorylation and degradation of the repressor IκBα [35–36], prompting the complete release of the canonical inflammatory transcription factor NF-κB (p65 subunit) from spatial inhibition and its rapid nuclear translocation [37–39], ultimately detonating a pro-inflammatory cytokine storm and extracellular matrix degradation in the large-vessel microenvironment at the nuclear transcriptional level.

For AAD patients, emergency surgery is the only treatment option. However, its high mortality, postoperative complication rates, and the lack of universal access to specialized cardiac centers highlight the necessity for new therapeutic strategies [1–3]. Therefore, addressing this clinical challenge by shifting focus toward targeting the GYS2-mediated immunometabolic network presents promising potential for non-surgical pharmacological intervention. Compared with the potential collapse of systemic host defense barriers caused by high-dose systemic glucocorticoids or traditional immunosuppressants [40], precise intervention targeting GYS2—the metabolic hub within macrophages—not only has the potential to intercept the continuous deterioration of aortic wall mechanical integrity at an ultra-early stage and reduce perioperative mortality but also offers a long-term active preventive strategy for high-risk populations with genetic predisposition or those in a pre-dissection state [41], thereby rewriting the traditional clinical paradigm that relies entirely on destructive surgical anatomical reconstruction.

Although this study constructed a rigorous causal mechanistic network, there are limitations. The primary consideration is that while the BAPN/Ang II mouse model used in this study highly recapitulates the AAD phenotype of medial elastic lamellar rupture and mechanical degeneration, there are undeniable species differences between rodents and humans regarding vascular anatomy, distribution of vasa vasorum, high-pressure hemodynamic shear stress, and basal immunometabolic substrate rates [42, 43]. Therefore, the safety margin, pharmacokinetics, and multi-organ tolerance of the GYS2-targeted blocking strategy in vivo must be handled with caution. Furthermore, while this study confirmed the superior vascular protective benefits of GYS2 genetic intervention, large-scale high-throughput screening of chemical small-molecule libraries has not yet been conducted; thus, significant work remains regarding the druggability evaluation of small-molecule targeted agents. Finally, although this work focused on macrophages as inflammatory “amplifiers,” clues from tissue immunofluorescence staining suggest that GYS2 also exhibits a certain degree of basal expression in vascular smooth muscle cells (VSMCs). As a core substrate metabolic enzyme, whether GYS2 participates in parallel in the pathological phenotypic dedifferentiation of VSMCs from a contractile to a synthetic phenotype, and whether a paracrine crosstalk [44–46] dependent on GYS2 metabolic derivatives exists between infiltrated macrophages and local VSMCs, remain to be clarified by further research.

In summary, this study not only provides a deep analysis of the molecular mechanisms underlying AAD progression but also offers a novel target and experimental support for clinical development of effective conservative therapeutic drugs. However, to translate these foundational discoveries into clinical applications, further exploration into the specific intervention timing for the downstream C5a/NF-κB signaling network across different pathological stages is required. Meanwhile, designing rigorous clinical cohorts and trials will be the essential pathway for finally verifying the translational feasibility of GYS2-targeted strategies.

## Acknowledgments

Not applicable

## Author Contributions

Jl, Ozh,Zl and Clw conceived and designed the study. Jl, Wwt,Mlk, Lmj, Jzs, Lyx, Xaq, Hxp and Xxm Performed the experimental procedures. Jl, Lmj, Hxp and Xxm Analyzed the data. Jl, Lyx, Xaq and Hxp Drafted the manuscript.

## Data availability

Availability of data and materials: The data that support the findings of this study are available from the corresponding author upon reasonable request.

